# Coupled mechanical mapping and interference contrast microscopy reveal viscoelastic and adhesion hallmarks of monocytes differentiation into macrophages

**DOI:** 10.1101/2022.11.29.518356

**Authors:** Mar Eroles, Javier Lopez-Alonso, Alexandre Ortega, Thomas Boudier, Khaldoun Gharzeddine, Frank Lafont, Clemens M. Franz, Arnaud Millet, Claire Valoteau, Felix Rico

**Affiliations:** Aix-Marseille University, INSERM, CNRS, LAI (U1067), Turing Centre for Living Systems, Marseille, France; Univ. Lille, CNRS, Inserm, CHU Lille, Institut Pasteur Lille, U1019 - UMR 9017 - CIIL - Center for Infection and Immunity of Lille, F-59000 Lille, France; Centuri, Turing Centre for Living Systems, Marseille, France; Univ.Grenoble Alpes, Inserm U1209, CNRS UMR5309, Institute for Advanced Biosciences, Team Mechanobiology, Immunity and Cancer, 38700 La Tronche, France; Department of Hepatogastroenterology, Centre Hospitalier Universitaire de Grenoble Alpes, 38700 La Tronche, France; WPI Nano Life Science Institute, Kanazawa University, Kanazawa, Japan

**Keywords:** cell mechanics, macrophages, viscoelasticity, atomic force microscopy, interference contrast microscopy, CD11b, holographic tomography

## Abstract

Monocytes in the blood torrent, when activated by pro-inflammatory signals, adhere to the vascular endothelium and migrate into the tissue for ultimately differentiate into macrophages. Mechanics and adhesion play a crucial role in macrophage functions, such as migration and phagocytosis. However, how monocytes change their adhesion and mechanical properties upon differentiation into macrophages is still not well understood.

In this work, we combined atomic force microscopy (AFM) viscoelastic mapping with interference contrast microscopy (ICM) to simultaneously probe, at the single-cell level, viscoelasticity and adhesion during monocyte differentiation. THP-1 monocytic cells were differentiated into macrophages through phorbol 12-myristate 13-acetate (PMA). Morphological quantification was achieved using holographic tomography imaging and the expression of integrin subunit CD11b was tracked as a marker of differentiation.

Holographic tomography proved to be a quantitative in vivo technique, revealing a dramatic increase in macrophage volume and surface area and two subpopulations, spread and round macrophages. AFM viscoelastic mapping revealed an increased stiffness and more solid-like behavior of differentiated macrophages, especially in the lamellipodia and microvilli regions. Differentiated cells revealed an important increase of the apparent Young’s modulus (E_0_) and a decrease of cell fluidity (*β*) on differentiated cells, which correlated with an increase in adhesion area. Macrophages with a spreading phenotype enhanced these changes. Remarkably, when adhesion was eliminated, differentiated macrophages remained stiffer and more solid-like than monocytes, suggesting a permanent reorganization of the cytoskeleton. We speculate that the more solid-like microvilli and lamellipodia might help macrophages to minimize energy dissipation during mechanosensitive activity, such as phagocytosis, making it more efficient. Our proposed approach revealed viscoelastic and adhesion hallmarks of monocyte differentiation that may be important for biological function.

## Introduction

Monocytes are circulating cells patrolling the vascular endothelium in search of external agents or inflammatory signals^1^. During inflammation, monocytes become adherent, able to trespass the vascular endothelium and the tissue to reach the site of injury or infection ^2,3^. To overcome external dangers, such as viruses or bacteria infections, monocytes differentiate into macrophages. Macrophages are highly specialized cells with a high phagocytic activity that once activated will be crucial for the inflammatory response and tissue healing^4–6^.

This process starts with the adhesion of the activated monocytes onto the vascular endothelium through specific interactions with adhesion molecules, like selectins and integrins ^1,7,8^. After activation and firm adhesion, monocytes trespass the endothelium deforming its cytoskeleton, forming pseudopodia and exerting traction forces to pull its body through the extracellular matrix^2,9^. Finally, monocytes differentiate into macrophages that are able to phagocytize external agents and start secreting inflammation mediators (pro-inflammatory and anti-inflammatory), until the tissue is back to its homeostatic state^10,11^.

The THP-1 monocytic cell line is known to differentiate into a macrophage phenotype through phorbol 12-myristate 13-acetate (PMA) and is a well-established biological model^12^. Exposure to the phorbol ester activates the PKC pathway, which stops proliferation and starts differentiation to macrophages^13,14^. This differentiation process has been extensively studied but quantification of relevant features has been elusive^15–19^. Biological changes like membrane antigens, secretory products (interleukins), and proto-oncogenes activation have been shown^16,20^. However, several morphological changes, like the increase in cell volume and cell diameter, have been difficult to quantify^12^. During differentiation, suspended monocytic cells become adherent, increasing the expression of adhesion molecules and leading to heterogeneous cell morphology with round, oval, or spherical shapes or with a spread membrane, stellate shapes, or ameboid shapes^12,18,19,21^. The expression of integrin CD11b is especially abundant in the podosomes of THP-1 PMA differentiated macrophages^22,23^. This integrin binds to multiple extracellular matrix (ECM) and vascular proteins, like fibrinogen and ICAM-1, and is commonly used as a marker for macrophage differentiation^8,24,25^. CD11b is also known to be a therapeutic target for other diseases like Lupus, and various types of cancer and has been used as a prognosis marker in myeloid leukemia^25–29^. Importantly, CD11b regulates macrophage polarization^30^.

Macrophage migration and phagocytic activity require adhesion molecules and the cytoskeleton to polarize towards the infection site. Integrins group around podosomes^22,23,31^, structures used by the cell to migrate and strongly adhere and which are directly linked to the cell cytoskeleton dynamics^32–34^. The state of the cell cytoskeleton directly modulates cell mechanics, cell shape and proportionate anchors for cell adhesion^35–38^. It has also been shown that adhesion structures are important for the development of stress fibers, cytoskeletal force transmission and cell shape^39–42^. Furthermore, it has been shown that leukocytes react to the stiffness of the substrate, for example, neutrophils display a spread morphology and become stiffer on stiff substrates and monocytes differentiate differently according to the substrate viscoelastic properties^43,44^. Moreover, monocyte firm adhesion to epithelial cells is mediated by integrins^45^ and monocyte adhesion is modulated by cell mechanics^46^. Previous studies have shown that, in the early stages (minutes) of the differentiation process, monocytes soften, increasing their adhesion capacity^47^, and that inflammatory response is sensitive to physical changes in the substrate^48^. Moreover, the mechanical properties of macrophages are involved in their activation state^49^, and cell mechanics allow distinguishing between pro-inflammatory (M1) and anti-inflammatory (M2) macrophages under inflammatory conditions^50,51^. Finally, during phagocytosis, macrophages stiffen, increase membrane and cortical tension, exert high traction forces and change the organization of the cytoskeleton^52–54^. Thus, cell mechanics and cell adhesion play an interconnected role in the physiological state of the leukocytes during the inflammatory response and are of vital importance in macrophage physiology. However, little is known about how adhesion and mechanics are related in fully differentiated macrophages. Therefore, it is important to measure cell mechanics and cell adhesion at the same time during the monocyte’s differentiation into macrophages.

Atomic force microscopy (AFM) is a robust technique providing highly quantitative data that allows the mechanical characterization of soft biological samples with nanometer resolution^55,56^. Coupling AFM with other optical techniques has been reported on several occasions^57–59^. For example, AFM for cell mechanics is commonly coupled to an inverted optical microscope for precise tip positioning over the sample. This gives access to using other techniques like immunofluorescence, confocal, total internal reflection fluorescence (TIRF) or stimulated emission depletion (STED) microscopy^60–64^. Interference contrast microscopy (ICM) imaging provides a measure of the distance of the cell’s membrane to the glass surface with nanometric precision and in a millisecond time scale, which makes it suitable to study adhesion in living cells^53,65–70^. Therefore, coupling AFM with ICM will allow to simultaneously determine viscoelasticity and adhesion on living cells.

In this study, we coupled AFM mechanical mapping to ICM to simultaneously determine cell viscoelasticity (apparent Young’s modulus and fluidity) and adhesion during monocyte differentiation into macrophages (Figure 1). The THP-1 monocytic cell line was differentiated into macrophages using PMA^12,71^. We first characterized cell morphology changes with label-free holographic tomography images, obtaining 3D holographic reconstructions of the cells^72^, and used flow cytometry to quantify the evolution of CD11b expression. This allowed initial characterization and quantitative classification of morphological populations. Differentiated macrophages increased their volume and showed two populations: spread and round. Viscoelastic maps showed that monocytes stiffened and became more solid-like (solidify) upon differentiation into macrophages, especially so in the spread areas. Adhesion correlated with this stiffening state. However, when adhesion molecules were cleaved and cells detached, resuspended macrophages remained stiffer and more solid-like than monocytes, suggesting permanent cytoskeleton reorganization.

**Figure 1.**
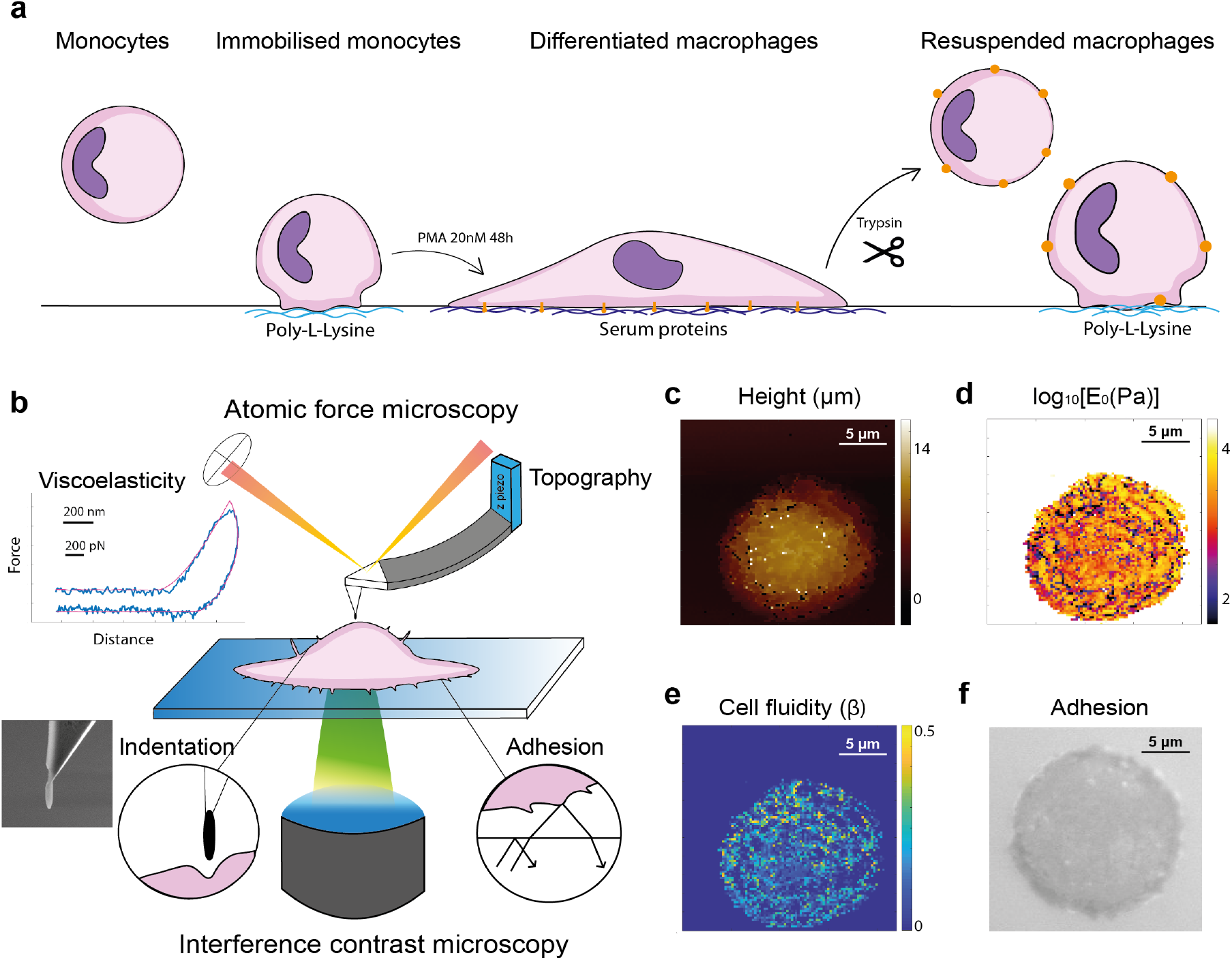
a. Experimental conditions to probe adhesion and viscoelasticity of monocytes and macrophages. a. THP-1 cells (monocytes) were grown in suspension and immobilized on poly-L-lysine (PLL) coated glass. THP-1 cells treated with 20nM PMA differentiated into macrophages after 48 hours and spontaneously adhered to the surface. Trypsin treatment cleaves proteins, like integrins, detaching macrophages that were immediately immobilized on PLL-coated surfaces. b. Coupled system of atomic force microscopy (AFM) and interference contrast microscopy (ICM). Optical interferences and AFM force curves allowed the formation of adhesion images and mechanical maps, respectively c. AFM mechanical mapping mode creates topography maps from the contact point detected in the force-indentation curves (inset in b). d. Apparent Young’s modulus (E_0_) and e. fluidity maps, were obtained from the viscoelastic fit (Eq. 1) to the force-indentation curves. f. Adhesion area of the very same cell obtained using ICM.

## Results

### CD11b expression

THP-1 cells were treated with PMA for 24, 48 and 72 hours (Supplementary Figure 1) and CD11b integrin cell surface expression was quantified over time using flow cytometry (Figure 2 a-b). PMA is known to induce the differentiation of THP-1 cells into macrophages, indicated by the upregulated expression of integrin subunit CD11b (ITGAM gene, forming Mac-1 with CD18)^12,15,18,19^. In agreement with previous reports, CD11b expression increased with time, reaching maximum levels after 48 hours of PMA treatment, and maintaining a similar level for 72 hours^19^. Thus, at 48h THP-1 cells were considered as differentiated into macrophages. PMA concentrations of 20 nM, 50 nM, 100 nM, and 200 nM were also tested obtaining similar results (Supplementary Figure 2). Thus, the minimum concentration needed for CD11b expression was used to induce differentiation, 20nM.

**Figure 2.**
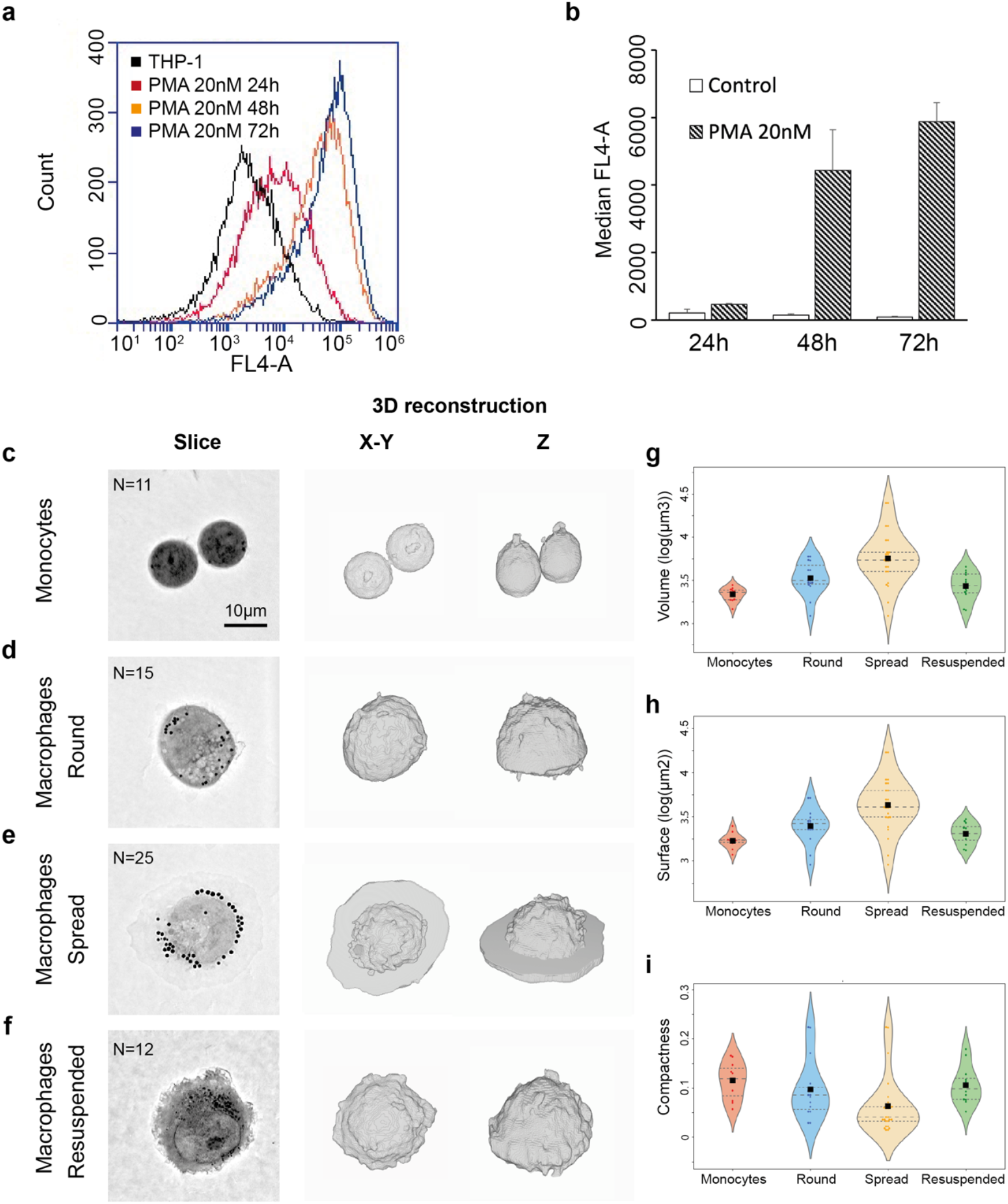
Molecular and morphological characterization of monocytes and macrophages. a-b Flow cytometry. a. Fluorescence intensity (FI) of CD11b obtained by flow cytometry on THP- 1 cells after 20nM PMA treatment at times 24h, 48h and 72h and control. b. CD11b mean +/- standard deviation of median FI (MFI) values from three flow cytometry experiments at 24h, 48h, 72h, for 20nM PMA treated cells and control. c-i. Tomography imaging of control and 20mM PMA treated THP-1 cells (48h). c-f. Image slices close to the substrate and 3D image reconstructions (from 96 slices) for monocytes, round, spread and resuspended macrophages. g-i. Volume, surface and compactness were determined after 3D single-cell reconstruction of tomographic images of monocytes and round, spread and resuspended macrophages (11, 15, 25, and 12 cells, respectively. Squares are mean, dashed lines are median, dotted lines 25% and 75% quartiles).

### Tomography imaging of cell membranes

To assess the morphological changes during THP-1 differentiation, individual cells were imaged by laser holographic tomography at 48h after 20nM PMA stimulation. This label-free technique generates 3D stacks and visualizes changes in refractive index throughout samples and is particularly suited for highlighting cellular membranes (Figure 2 c-f). At 48h, macrophages presented two phenotypes: 1) a prevalent round morphology (Figure 2 d), and 2) a spread morphology with roughly circular areas on the substrate (Figure 2 e). Similar changes in morphology have been identified before as a signature of differentiation. We, therefore, analyzed both morphological phenotypes separately and used image analysis to quantify the volume, surface area and compactness of the cells before and after differentiation (Figure 2 g-i). Differentiated macrophages displayed lower compactness, larger volume, and larger surface area than monocytes. The spread phenotype had the lowest compactness, as expected given the less spherical shape, and the largest volume and surface area (both ~2.6 times larger than monocytes). In addition, tomography imaging revealed that all differentiated macrophages contained more and larger cytoplasmic vesicles^12,73^. To assess the effect of substrate adhesion on cell morphology, macrophages were resuspended using trypsin, immobilized on PLL-coated bottom dishes and immediately imaged. Resuspended cells did not spread and presented reduced volume and surface area and slightly higher compactness than the round phenotype (Figure 2 g-i). Therefore, this technique allowed us to classify cells into four states: monocytes, round and spread macrophages and resuspended macrophages.

### Mechanical properties and topography

To determine how the viscoelasticity changed among the different groups described, AFM was used to obtain mechanical maps on individual cells (30×30μm with 64×64 pixels, 468 nm/pixel), containing 4096 force-distance curves. The apparent Young’s modulus (E_0_) and the cell fluidity (*β*) were determined by fitting a parabolic viscoelastic contact model to each force-indentation curve (Equation 1). Thus, maps revealed topography, apparent YM and cell fluidity. (Figure 3 a-c). Topography was extracted from the contact point determined by the fit, reflecting the undeformed cell surface. Topography maps allowed us to discern between the different cell morphologies, spread macrophages showing a spread membrane of ~400 nm thickness (Figure 3 a).

**Figure 3.**
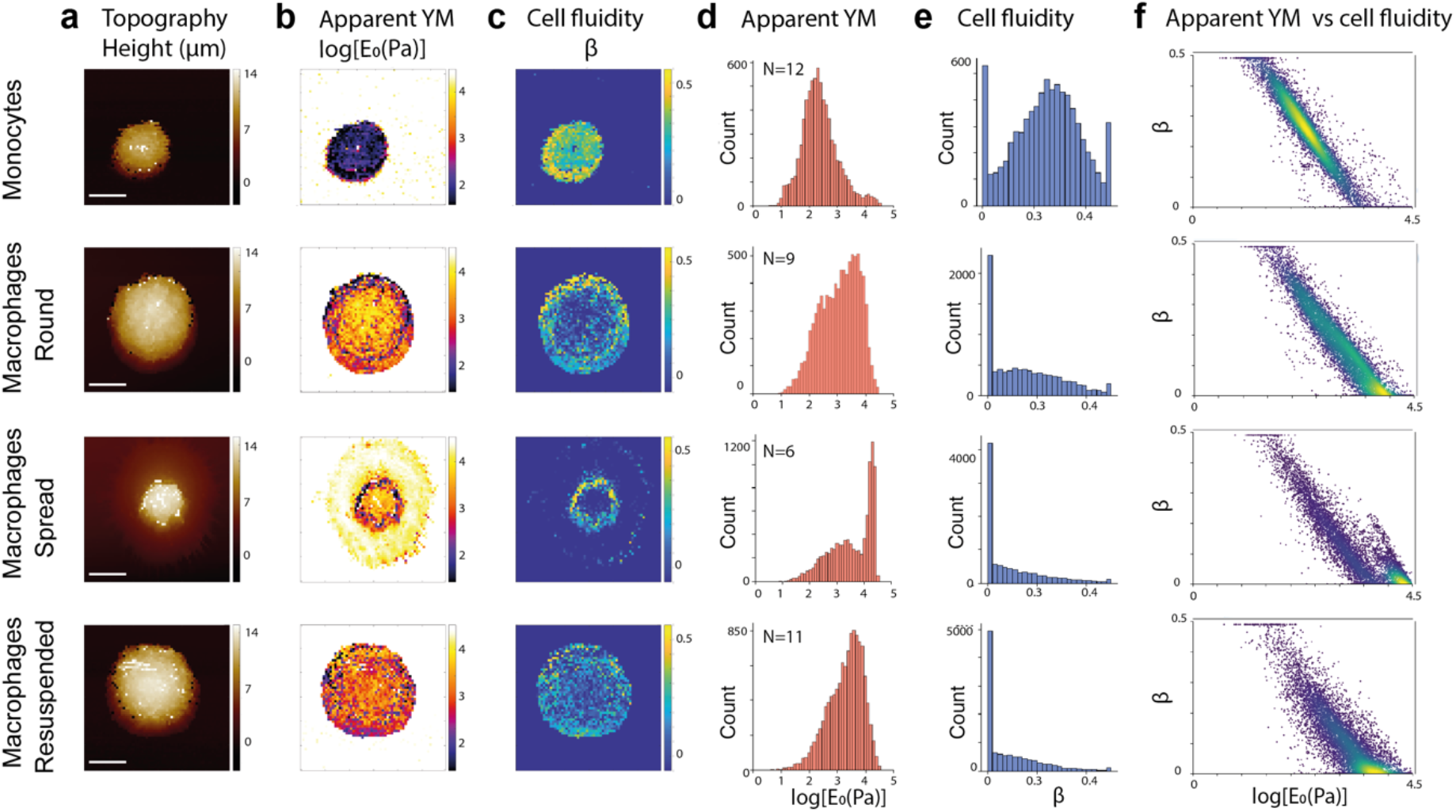
Atomic force microscopy topography and viscoelastic maps. 30×30 μm, 64×64 px, 4096 F-d curves per map at 468 nm/pixel. a. Topographical maps extracted from the contact point of the F-d curves (scale bar 10μm). b. Apparent YM maps (E_0_), color scale: 1.7-4.47 log[Pa], with saturated values on the glass surface. c. Cell fluidity maps (*β*), color scale: 0-0.5. d. Apparent YM (E_0_) distribution in log scale. e. Cell fluidity (*β*) histogram. f. Cell fluidity (*β*) versus log[*E*_0_]. Monocytes N=12, round macrophages N=9, spread macrophages N=6, resuspended macrophages N=11. The number of curves for d-f was between 8000 and 10000 per condition studied after filtering out the background curves.

From apparent YM (E0) maps, macrophages appeared to be on average stiffer than monocytes, regardless of the cell state. Apparent YM (E0) maps allowed us to assess the distribution of stiffnesses across the cell (Figure 3 b). On spread macrophages, the membrane extension appeared to be ~10-fold stiffer than the body of the cell. Also shown in Figure 3 b, the body center of round and spread macrophages was stiffer than the body periphery, while the values were more homogeneous in resuspended macrophages. Protrusions and microvilli appeared to be stiffer than the average body. Histograms of log(E_0_) pooling all maps were generated, further evidencing the differences between cell states (Figure 3 d). Monocytes showed a distribution of log[E_0_] values with mean±SD of 2.38±0.63 log[Pa] (~240 Pa). For macrophages, the mean±SD appeared at 3.08±0.69 log[Pa] (~1200 Pa) (round) and 3.53±0.78 log[Pa] (~3390 Pa) (spread). Interestingly, spread macrophages presented a histogram with clear two peaks, at 3.2 log[Pa] (~1580 Pa) and 4.3 log[Pa] (~20 kPa), corresponding to the cell body and the spread regions, respectively, as observed in the mechanical maps (Figure 3 b). Interestingly, resuspended macrophages showed a mean±SD of 3.20±0.61 log[Pa] (~1580 Pa), slightly stiffer than round macrophages.

Fluidity (*β*) maps revealed the distribution of solid- (*β*~0) and fluid-like (*β*~1) regions across the cell, generally revealing low *β* values at high E_0_ regions (Figure 3 c). As expected for a stiffer phenotype, cell fluidity decreased in macrophages, suggesting a more solid-like response than monocytes. Solid-like areas, with *β*~0, were more frequent in the cell centre and borders, corresponding to the areas with microvilli and filopodia, respectively. Histograms of *β* values were generated pooling together all maps (Figure 3 e), revealing mean±SD of 0.24±0.12 for monocytes, 0.15±0.13 for round macrophages and 0.09±0.12 for spread macrophages. Interestingly, resuspended macrophages had a mean±SD beta of 0.08±0.10, similar to spread macrophages. Monocytes showed a symmetric and centered distribution, with two peaks at the limits 0 and 0.5. In contrast, macrophages showed distributions with a peak near or coincident with the lower limit of 0 and a long tail towards higher *β.* To notice, *β* values equal to zero were very frequent, reflecting a perfectly elastic response.

In all states, a linear trend was observed between *β* and log[E_0_] (Figure 3 f). However, from monocytes to macrophages, the peak was shifted towards lower *β* and higher log[E_0_]. In the case of spread macrophages, two clear populations were observed, corresponding to the body and spread areas (Figure 3 f).

### Cell adhesion by ICM

The coupled AFM/ICM system allowed to directly quantify the adhesion of cells to the substrate (Figure 4) on the very same cells mapped by AFM. This allowed direct correlation between adhesion and viscoelastic parameters. The contrast in ICM image pixels values is related to the distance between the cell membrane and the glass surface, following a damped sinusoidal relationship as the distance increases, with a wavelength of ~200 nm (half the wavelength of incident green light, *λ,* divided by the refraction index of the medium n=1.33 for water ^65^). The periphery of cells was brighter than the cell center and spread regions (thinner than *λ*/2) appeared darker. Thus, we assumed that darker regions (lower pixels values) implied shorter distances, i.e. stronger adhesion, as suggested before^68^. Therefore, pixel values provide a semi-quantitative measure of cell adhesion strength. The ICM images were processed as explained in the methods and the distribution of pixels values from the cell area was plotted as averaged histograms (Figure 4 a-d). Macrophages showed lower pixel values, suggesting that macrophages of any state had stronger adhesion compared to monocytes. ICM images on macrophages revealed more variability in the grey values, often showing small, circular dark areas, resembling podosome-like structures, and circular ruffles suggesting actin bundles observed before^18,22,31^. Adhesion decreased in resuspended macrophages, whose intensity histogram showed a peak in a similar range as monocytes, with a second peak at lower values suggesting punctual strong adhesion areas.

**Figure 4.**
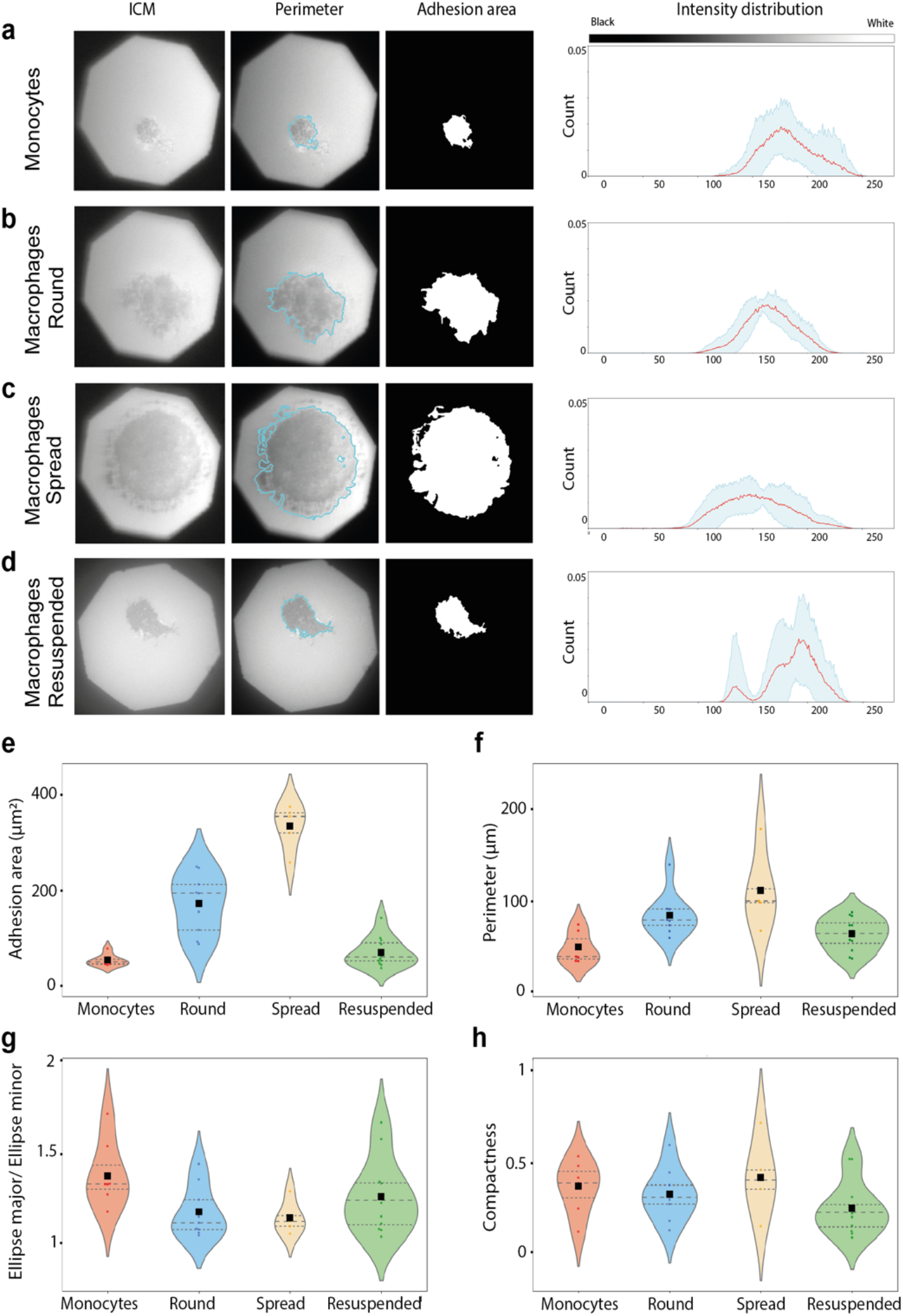
Interference contrast microscopy (ICM) image processing. a-d. ICM images (left) for monocytes and round, spread and resuspended macrophages. ICM images were processed to extract the mask and quantify the adhesion area (right) and perimeter of each cell (centre). The intensity values inside the mask for all cells at each state were pooled and an average histogram (red line) was generated, with values ranging from 0 (black) and 255 (white) (blue shading represents the standard deviation). e. Adhesion area (μm^2^). f. Cells perimeter (μm). g. Ratio ellipse major axis/ellipse minor axis. h. Compactness. Monocytes N=9, round macrophages N=9, spread macrophages N=6, resuspended macrophages N=11.

ICM images were processed as described in the methods to quantify the adhesion area (μm^2^), perimeter (μm), ellipse major/minor ratio, and compactness (Figure 4 e-g). Macrophages showed larger adhesion area than monocytes, especially in the spread phenotype (4 and 6-fold larger for round and spread, respectively), recovering monocyte values when resuspended (Figure 4 e), in accordance with tomographic imaging results for surface and volume (Figure 4 g-h). The perimeter followed the same but less pronounced trend (Figure 4 f). The ratio between the ellipse major and minor diameters showed that spread macrophages had rounder adhesion areas (Figure 4 g). The compactness of the adhesion area did not vary importantly, although the lowest value was found in resuspended macrophages suggesting rougher perimeters (Figure 4 h).

To summarize, macrophages appeared stiffer and more solid-like than monocytes in all conditions. These results were more pronounced on spread macrophages. Overall, stiffness and solidity correlated with adhesion, the more adhesive cells being stiffer and more solidlike. However, resuspended macrophages did not recover monocyte viscoelasticity, maintaining the solid-like behavior of spread macrophages with the apparent Young’s modulus of round macrophages.

## Discussion

Tomography imaging revealed the cell surface *in vivo,* without labeling and with nanometric resolution (183 nm in xy, 312 nm in z), while 3D reconstruction allowed us to determine the morphology of cells, providing a quantitative measure of the different cell phenotypes (Figure 2 g-i). Compared to monocytes, differentiated macrophages, regardless of the condition, increased volume and surface, and reduced compactness. As reported before, macrophages often presented round phenotypes and, less frequently, membrane spreading (Figure 2 d and e, respectively)^12,18^. We named these two populations round and spread macrophages. This first visual division corresponds to a 1.7-fold increase in surface and volume and a 1.5-fold decrease in compactness of spread macrophages compared to round ones. Resuspended macrophages presented similar compactness, and slightly smaller volume and surface than round macrophages (Figure 2 g-i). Therefore, tomography imaging allows the establishment of cell populations based on quantitative morphological features of unlabeled cells in-vivo. Recent work on blood samples from COVID-19 patients revealed increased size and volume of monocytes (~1.2-fold) due to dysregulated inflammatory response or cytokine storm syndrome^74^, which, according to our quantification, may be an early signature of differentiation. In this direction, COVID-19 patients’ blood samples have also reported monocytes expressing macrophage markers like CD80 and CD206^75^.

Tomography imaging also allows visualization of internal membranes, such as vesicles (Figure 2 c-f). THP-1 cells after differentiation showed an increased number of small, dark spheres, resembling vesicles, characteristic of macrophages^73^. Lysosomal vesicles are related to the antimicrobial activity of macrophages and their content is released in case of pathogenicity. In the resuspended phenotype, the number of vesicles diminished, suggesting release during resuspension, which may explain the reduced volume and surface area. Therefore, vesicle number may be used as an additional quantitative marker of monocyte differentiation.

AFM and ICM were combined to simultaneously obtain topography, viscoelasticity and adhesion maps on the very same cells. This coupling approach has been used before to characterize hyaluronon brushes and living cells^57,76^, but not to correlate adhesion and mechanics on cells. A possible improvement of the current setup would involve adding a lambda quarter waveplate between the objective and the sample and using different illumination wavelengths. This would help increase the signal-to-noise ratio and allow direct quantification of the substrate-membrane distance^65,70^.

To obtain mechanical maps of living monocytes and macrophages, we used long and relatively sharp AFM tips. Long tips of ~20 μm were necessary to allow mapping of 10-14μm height cells without touching the cell body with the cantilever arm. Moreover, long tips allowed us to obtain force curves at a relatively high velocity (~200 μm/s) with minimal contribution of the substrate to viscous drag forces on the cantilever^77^. Finally, relatively sharp tips (~30 nm) provide relatively high-resolution maps, revealing the structure and mechanics of the cell surface with submicrometer resolution. While we were able to obtain good quality maps on living cells at 234 nm/pixel (Figure 1), the long time acquisition (30 min) limited the viability of the cells and thus, we opted to use a lower resolution (468 nm/pixel). Nevertheless, the maps allow visualization of subcellular structures, such as filopodia, and quantify their mechanical properties (Figure 3). This was important to detect the contribution of cell stiffness and fluidity of the different regions across the cell surface.

Compared to monocytes, maps revealed stiffening of macrophages in all states, as showed the geometric mean of the apparent YM (*E*_0_, Figure 3). Round macrophages (1202 Pa on average, beta=0.15) were 5 times stiffer and considerably less viscous than monocytes (240 Pa, beta=0.24). Remarkably, spread macrophages were ~14-fold stiffer than monocytes and even more solid-like than round macrophages (3388 Pa, beta=0.09). This dramatic increase in E_0_ was due to its bimodal distribution, corresponding to the cell body and spread area, with peaks at 1584 Pa and 20 kPa, respectively. The first peak of the apparent YM distribution corresponded to the cell body, slightly stiffer than round macrophages, being this region less viscous, suggesting that the spreading may induce tension or prestress in the cell cortex^78,79^. The second peak of the YM distribution correlated with the spread region and was ~80 times stiffer than monocytes and almost perfectly elastic (beta~0). Measurements in thin spread regions (400 nm thick) may appear stiffer due to the hard bottom effect. However, for an average indentation of around 200 nm (Supplementary Figure 5) and given the relatively sharp tip (~30 nm), we evaluated this overestimation to be less than 20%, another advantage of using relatively sharp probes. Therefore, spread regions were remarkably stiffer and almost perfectly solid-like. This may be important for phagocytic activity.

In addition, the resolution of the maps allowed us to discern stiffer and solid-like regions coincident with protrusions, resembling microvilli. A large proportion of cell surface appeared purely elastic, in particular at lamellipodia and microvilli. Macrophages are highly mechanosensitive cells^5,49,80–83^, a more solid-like response of lamellipodia and the apical part of microvilli would result in higher mechanosensitivity and more efficient downstream transmission of the mechanical stresses. Thus, we conclude that monocytes differentiate into macrophages by developing high mechanosensitive solid-like regions, lamellipodia (membrane spreading) and microvilli and protrusions (macrophage central body). These solidified structures may render mechanosensing more efficient, minimizing the dissipation of energy due to viscous effects. A similar purely elastic response has been recently reported on the periphery/boundaries of various adherent cell lines^84^, suggesting solidification as a possible fingerprint of mechanosensing during cell spreading and migration.

Spread macrophages extended the membrane, modulating both cell stiffness and adhesion to the substrate, resembling what has been termed frustrated phagocytosis^18,53^. As previously reported contact activated neutrophils on stiff surfaces changed to a spread morphology with a stiffer and more solid-like mechanical response, while monocytes differentiate aberrantly on stiff substrates^43,44^. The stiffening of leukocytes during inflammation has been reported and hypothesized to be a fingerprint of leukocyte activation, inducing adhesion and extravasation^74,85,86^. It has also been described in several works that macrophage activation state in M1 or M2 phenotypes change their mechanical properties^49,81^. In apparent contradiction, THP-1 cells treated with PMA have been reported to soften in the short term (~30min), enhancing cell adhesion^47^. In contrast, after long-term stimulation (48h), we observed stiffer macrophages, likely due to cytoskeleton reorganization, accumulation of actin filaments in the cell edge and podosome formation^18,22,31^. This suggests that PMA-induced short-time softening (minutes) helps monocytes to adhere to the substrate, consequently inducing cytoskeleton remodeling, stiffening and solidification at longer times. These changes may allow macrophages to migrate and phagocyte external agents more efficiently^12,14,19^. Interestingly, when macrophage adhesion was suppressed through trypsinization, macrophages still remained 6.6-fold stiffer (1585 Pa) than monocytes, probably due to a permanent reorganization of the cytoskeleton, likely of actin and associated proteins^23,87^. Taken together, these results suggest that macrophage stiffening is due both to cell adhesion-mediated tension and cytoskeleton remodeling.

ICM allowed us to evaluate the changes in adhesion of the different cells and correlate them later with the mechanical state (Figure 4). Adhesion area and perimeter increased during differentiation of monocytes into macrophages (Figure 4 e-f). This is partly due to the increase in the size of the cells observed by tomography 3D reconstructions images (Figure 2 c-f). However, the size increase was up to 2.6-fold in 3D (for spread, 1.5 for round), while the increase in adhesion area was between 3.2- and 6.6-fold, suggesting an important amount of excess membrane in macrophages, allowing phagocytosis of large external agents. In addition, the adhesion areas appeared to be circular for macrophages (Figure 4 g). These changes were more accentuated in the spread macrophages. Interestingly, the increase in adhesion area and perimeter correlated with the increase in adhesion strength as observed from the darker ICM intensity distributions, again suggesting frustrated phagocytosis (Figure 4 a-d). Given that CD11b expression increases dramatically at 48h, it seems reasonable that the stronger and larger adhesion was mediated substantially by integrin Mac-1, formed by subunits CD11b and CD18, likely binding to serum proteins like fibrinogen adsorbed to the glass surface^18,19,21^. When macrophages were resuspended, the adhesion returned to monocyte levels. Some ICM images of macrophages only presented darker spots, which could correlate to strong adhesion areas related to podosomes observed before in these cells^18,22,31,68^.

The relationship between adhesion and mechanics in macrophages has been suggested before. The adhesiveness of THP-1 cells to various ligands has been measured by optical tweezers, showing an increase in binding forces after 48h PMA differentiation that correlated with higher traction forces exerted by cells^88^. In addition, stiffening of extracellular matrix due to crosslinking increased adhesiveness in THP-1 PMA-differentiated cells^9^. Quantification of the apparent Young’s modulus, fluidity and adhesion area on the very same cells allowed us to directly compare the average parameters against each other. Overall, monocyte differentiation revealed a correlation between adhesion and viscoelastic parameters (Figure 5 a-b). Monocytes differentiated into macrophages stiffened, solidified and showed larger adhesion area. However, within populations, larger adhesion did not always result in stiffer and more solid-like cells. This is clearly observed in spread macrophages, in which even a negative correction of log[E_0_] vs adhesion is observed (Figure 5 a). Thus, changes in cell mechanics may be due solely to integrin expression, as observed recently on melanoma cells^89^, although may be reinforced during the initial steps of firm adhesion. After this, a possible mechanical upper bound may limit the relationship between adhesion expansion and stiffness, as suggested from neutrophil phagocytosis^90^. In accordance, when adhesion was suppressed in resuspended macrophages, the viscoelastic parameter values did not recover monocyte levels (Figure 5 a-b). This suggests that adhesion, stiffening and solidification progress together during differentiation but that may become independent of each other once differentiation is reached.

**Figure 5.**
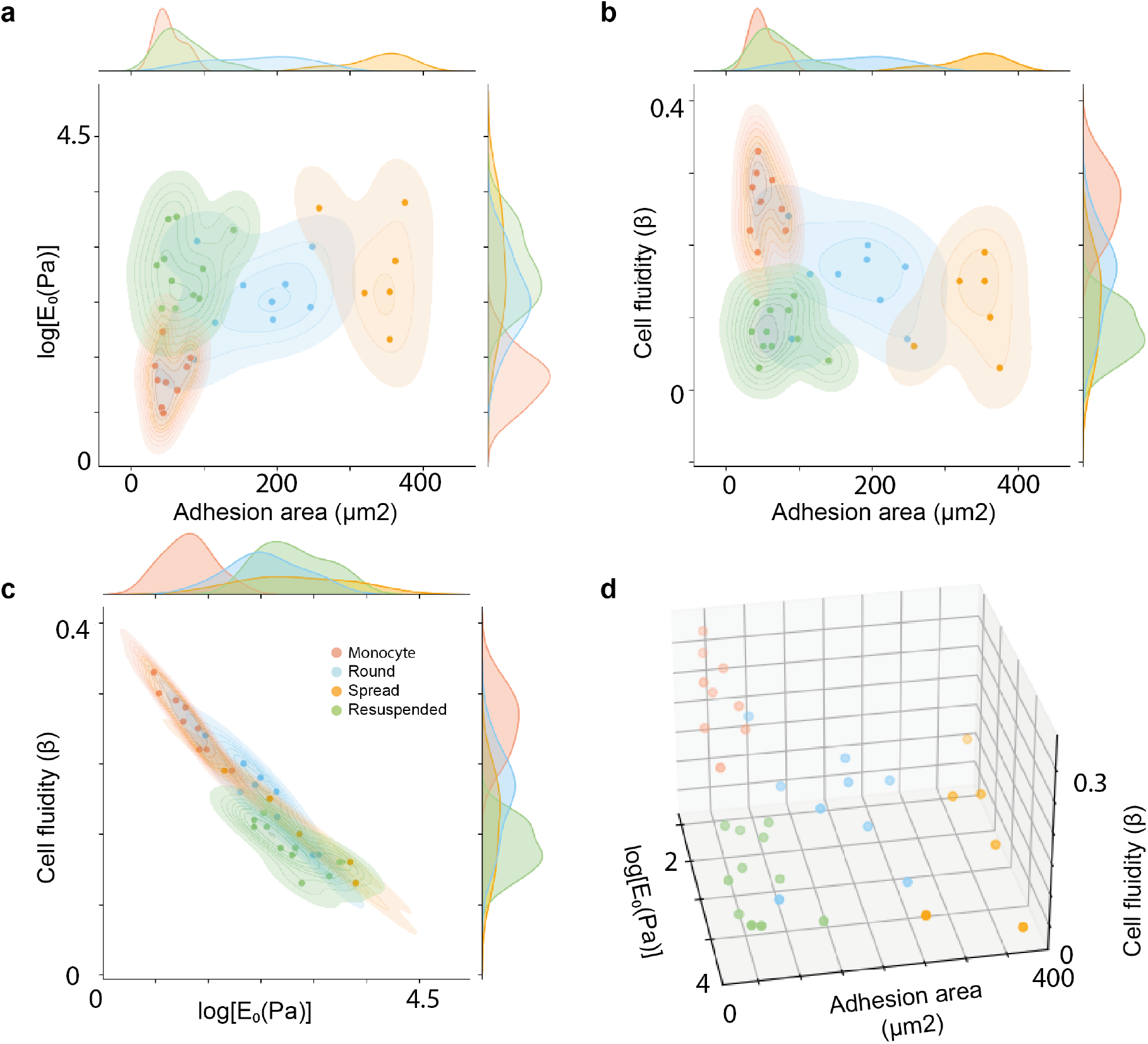
Viscoelasticity and cell adhesion area comparison for single cell analysis (each symbol corresponding to a single cell). a. Apparent YM (Log_10_[E_0_]) versus adhesion area (μm^2^). b. Cell fluidity (*β*) versus adhesion area (μm^2^). c. Cell fluidity (*β*) versus apparent YM (Log_10_[E_0_]). d. Cell fluidity (*β*), apparent YM (Log_10_[E_0_]) and adhesion area. Different colors correspond to different cell conditions.

It is interesting to notice that, on average, monocytes, round and spread macrophages all fell within the same trend line of beta vs log[E_0_], while resuspended macrophages slightly deviated from this trend. Interestingly, using the three main parameters as dimensions of a phase space, we observed that the four cell phenotypes clustered around different regions (Figure 5 d).

The correlation between adhesion and viscoelasticity and the increased expression of CD11b suggests that integrin expression may be the triggering factor to induce stiffening and solidification. However, mechanical hallmarks of macrophages remain independent of adhesion once differentiation was reached. CD11b has been reported as a key biomarker in immune cells for the progression of diseases like cancer and autoimmunity^25,28–30,91–93^. Given the growing evidence that cell elasticity and fluidity suffer alterations in disease, adhesion and mechanics of immune cells may appear as relevant biomarkers^26,94–103^.

## Conclusions

In conclusion, we applied holographic tomography imaging as a quantitative, non-invasive and label-free technique uncovering round and spread macrophage phenotypes with larger volume and surface area than monocytes. Coupling of AFM mechanical mapping with ICM allowed us to quantify the adhesion and viscoelasticity on the very same cells revealing viscoelastic hallmarks of monocyte differentiation into macrophages that correlated with adhesion. Compared to monocytes, macrophages showed larger adhesion, higher apparent YM (E_0_) and decreased cell fluidity (*β*), particularly in lamellipodia and microvilli. Mechanical changes may be a consequence of adhesion to the surface through a reinforcement mechanism, as shown in the spread macrophages whose population stiffening almost tripled compared to the round phenotype, mainly due to the spread regions. However, this correlation fails within spread macrophages when using average cell values. Recent work has reported stiffening and solidification of neutrophils and macrophages during phagocytosis^90^, which might be frustrated in the spreading phenotype^53^. Stiffening and solidification may help force transmission. Taken together, our results support the idea that stiffening, solidification and size change of monocytes during differentiation into macrophages are biologically important. We propose that larger, stiffer, and solidified macrophages may be more efficient during mechanosensitive activities, such as migration and phagocytosis^104,105^.

## Methods

### Cell culture

The THP-1 cell line was obtained from ATCC (American Type Culture Collection, TIB-202) and cultured in RPMI-1640 media (Gibco, Thermofisher, Ref.11875093) supplemented with 1% sodium pyruvate (Gibco, Thermofisher, Ref. 11360070), 1% of MEM NEAA (Gibco, Thermofisher, Ref. 10370047), 1% Penicillin-streptomycin (Gibco, Thermofisher, Ref. 15140148), HEPES 10mM (Gibco, Thermofisher, Ref.15630080) and 10% fetal bovine serum (FBS). Cells were maintained at 37º, 5% CO2 and 95% humidity. The cells were kept between a concentration of 2-8 x 10^5^ cells/mL, splitting them every 2-3 days. Cells were discarded after 20 passages to avoid any drifting in the phenotype. All % values are v/v %. Cells were routinely tested for mycoplasma contamination.

### Cell differentiation

THP-1 cells were cultured in the cell culture media described before with 20nM phorbol 12- myristate 13-acetate (PMA) (Sigma Aldrich, Ref. P1585) for 48h in glass-bottom Petri dishes. After 48h the cells were washed once with fresh cell culture media, without PMA, prewarmed at 37°C.

### Atomic force microscopy measurements

Cultured cells’ viscoelastic properties and topography were measured with AFM-force mapping mode in a Nanowizard 4 (Bruker-JPK) using pre-calibrated PFQNM-LC-cal cantilevers (Bruker). The force set point was set at 0.8 nN, speed at 200 μm/s and the range at 15μm in Z. The first experiments were conducted at high resolution (234 nm/px) and then the resolution was set to 64×64px in a 30μm square area (468 nm/px). We applied the SNAP approach to calibrate the inverse of the optical lever sensitivity (invOLS) from the thermal spectra in liquid^106^. For that, we used the pre-calibrated spring constant of the cantilevers and the correction factors described in Rodríguez-Ramos 2021^107^.

The viscous drag coefficient of the cantilever (b=0.7 pN·s/μm) was determined as the difference between approach and retract forces before contact from force curves obtained on top of the cells. For the monocyte and macrophage measurements, the cells in suspension were immobilized with a poly-L-lysin 0.01% solution coating of high molecular weight (>80kDa), (Sigma Aldrich Ref.25988-63-0). For the adhesion control, THP-1 differentiated cells were trypsinized for 3-5 minutes at 37°C, centrifuged at 1200rpm for 10 min and seeded with fresh culture media without PMA in a glass-bottom petri dish coated with poly-L-lysine. The measurements of all conditions were taken in the culture media without PMA, at room temperature and 20mM Hepes in a window of time of 2 hours at room temperature.

### Viscoelastic properties

The apparent Young’s modulus (E_0_) and cell fluidity (*β*) were determined using a parabolic contact model assuming a viscoelastic sample following power law rheology of time-dependent Young’s modulus *E*(*t*)=*E*_0_(*t*/*t*_0_)*β* ^108^, [Lacaria 2022 in preparation]. Briefly, the approach and retract force versus time traces (*F_a_*(*t*) and *F_r_*(*t*), respectively) were fitted simultaneously using the equations below.

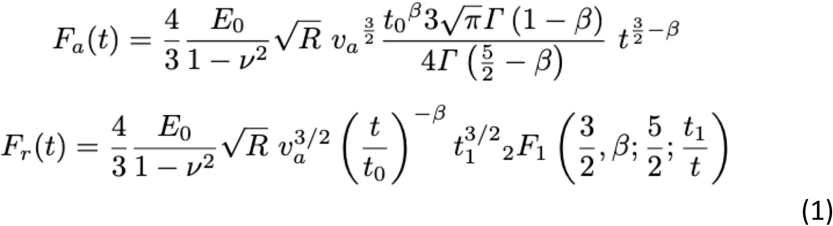

[Lacaria 2022 in preparation] where 2*F*_1_ is the ordinary hypergeometric function, *v*=0.5 is the Possion ratio, *v*_a_ and *v*_t_ are the approach and retract velocities, respectively, *t*_0_=1s and *t*_1_ is the time point from the retract curve at which the area of contact equals the area of contact of the approach and is found to be

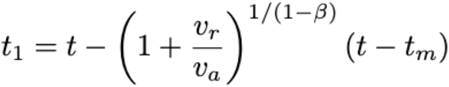

where t_m_ is the time at maximum indentation. The viscous drag force was added to the approach and retract curves from the viscous drag factor times the velocity calculated at each time point: b·v_a_(t) and -b·v_r_(t), respectively.

### Interference contrast microscopy (ICM)

The ICM observations were done *in situ* using the optical microscope Ti-Eclipse (Nikon, Japan) coupled to the AFM. The images were acquired using an epi-illumination (light source LHS- H100C-1, Nikon, Japan) featuring a cube equipped with a polarizer, an analyzer and a semi-reflective mirror (Nikon, Japan), a 60x water immersion objective (NA 1.2, Nikon, Japan) and a CCD camera (model C8484-03G01, Hamamatsu, Japan). The AFM laser was shut down and the aperture diaphragm was closed to illuminate only the area of interest. The images were acquired using the HCImage software (provided by Hamamatsu), automatically adjusting the acquisition time.

### ICM Image processing

The images were processed using a self-written Python 3.8.8 code. Briefly, the images were first segmented to select only the pixels corresponding to the inner part of the aperture diaphragm, using the unsupervised learning method Kmeans (from the sklearn package). A median blur was then applied to remove the highly-contrasted points (corresponding to impurities in the optical path and/or on the Petri dish surface). An estimation of the background was performed using the rolling ball method (with skimage restoration rolling ball function) and subtracted to the raw image to homogenize the brightness over the full field of view. The remaining noise was removed using Chambolle projection algorithm (with skimage restoration denoise-tv-chambolle function). To reveal the contour of the contact area, a map of the local variations was generated using a circular kernel and a mean-C local thresholding (also called adaptive thresholding) was applied. When the cell was centered in the shutter, the contour was closed using the corresponding function from the skimage module, and a flood fill was realized. When the cell ICM pattern was near the border of the diaphragm, to avoid the shutter to be detected as part of the cell, an eroded version of the inner shutter mask was applied to the image. In both cases, only the biggest filled area (corresponding to the cell) was kept, the others (corresponding to impurities) were discarded. This area is slightly overestimated due to the size of the local variance kernel. To refine the area estimation, this overestimated area was used as a mask and applied to the raw image. Any pixel whose value is greater than the mean value of all pixels inside this mask was removed (considered as the background). A graphical user interface was developed (with pygame package) for the user to decide whether the detected surface is correct or not, allowing the user to draw on the image to discard or add parts. Finally, multiple parameters, such as the area, perimeter or compactness, are measured using the skimage measure regionprop method.

### Holographic tomography and image processing

Cells were seeded in glass-bottom Petri dishes using the same conditions as for AFM-force mapping and imaged by holographic tomography^72^ using a 3D Explorer (Nanolive). Live cells were imaged in RPMI 1640 medium with a normalization refractive index set to 1.338. The collected image stacks were analyzed in ImageJ and TAPAS to extract single-cell volume and surface area^109,110^. Two distinct cell regions were extracted separately: the main part corresponding to the highly-contrasted central part of the cell, and the remaining part corresponding to thin membrane extensions at the cell perimeter. The central cell region was detected using a classical segmentation protocol. For that, stacks were filtered with a 3D median filter with radii 4×4×2, followed by automatic thresholding (using the triangle algorithm) and a “fill holes” procedure. The weakly-contrasted thin membrane boundaries were manually delineated. Finally, the individual values computed for the central and peripheral regions were combined to obtain total single cell volume and surface areas.

### Flow cytometry

THP-1 cells were cultured in wells as detailed in the cell culture methods and analyzed using a flow cytometer (BD Accuri C6 flow cytometer, BD Biosciences). Differentiated cells were detached using Enzyme Express (1X) TrypLE™ (Gibco, Ref.12605010), to preserve the adhesion proteins on the surface, and collected in FACS tubes for analysis. Cells were stained with CD11b Antibody, anti-human, REAfinity™ (Miltenyi Biotec, Ref.130-110-554) and REA control (I)-APC human for the control of unspecific binding (Miltenyi Biotec, Ref. 130-104- 615). At least 10000 events were gated according to the FSC/SSC dot plot with the exclusion of dead cells.

## Acknowledgements

The project was supported by the European Union’s Horizon 2020 research and innovation programmes No 812772 (project Phys2BioMed, Marie Skłodowska-Curie grant), the Japanese Society for Promotion of Science (JSPS) short-term fellowship, the European Research Council (ERC) under the European Union’s Horizon 2020 research and innovation programme (grant agreement No 772257), and the ATIP Avenir with financial support from ITMO Cancer of Aviesan on funds Cancer 2021 administered by Inserm. The project leading to this publication has received funding from France 2030, the French Government program managed by the French National Research Agency (ANR-16-CONV-0001) and from Excellence Initiative of Aix-Marseille University - A*MIDEX. We thank Dr Millet and his laboratory for providing the low-passage THP-1 cells from ATCC (American Type Culture Collection, TIB-202). We also want to acknowledge the support of WPI NanoLSI and Turing Centre for Living Systems (Centuri).

## Notes

### Competing Interest Statement

The authors have declared no competing interest.

