## Supplementary material for "Coupled mechanical mapping and interference contrast microscopy reveal viscoelastic and adhesion hallmarks of monocytes differentiation into macrophages": SupplementalFile.pdf

SUPPLEMENTARY INFORMATION

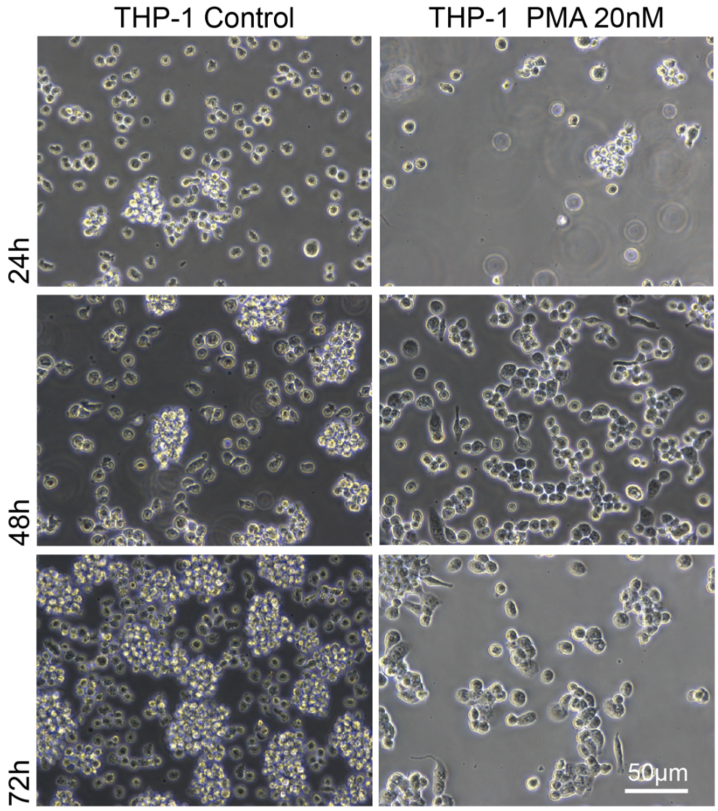


Supplementary Figure 1. PMA differentiation of THP-1 cells over time. a. Phase contrast images of THP-1 cells control and 20nM PMA treated cells at 24h, 48h and 72h.

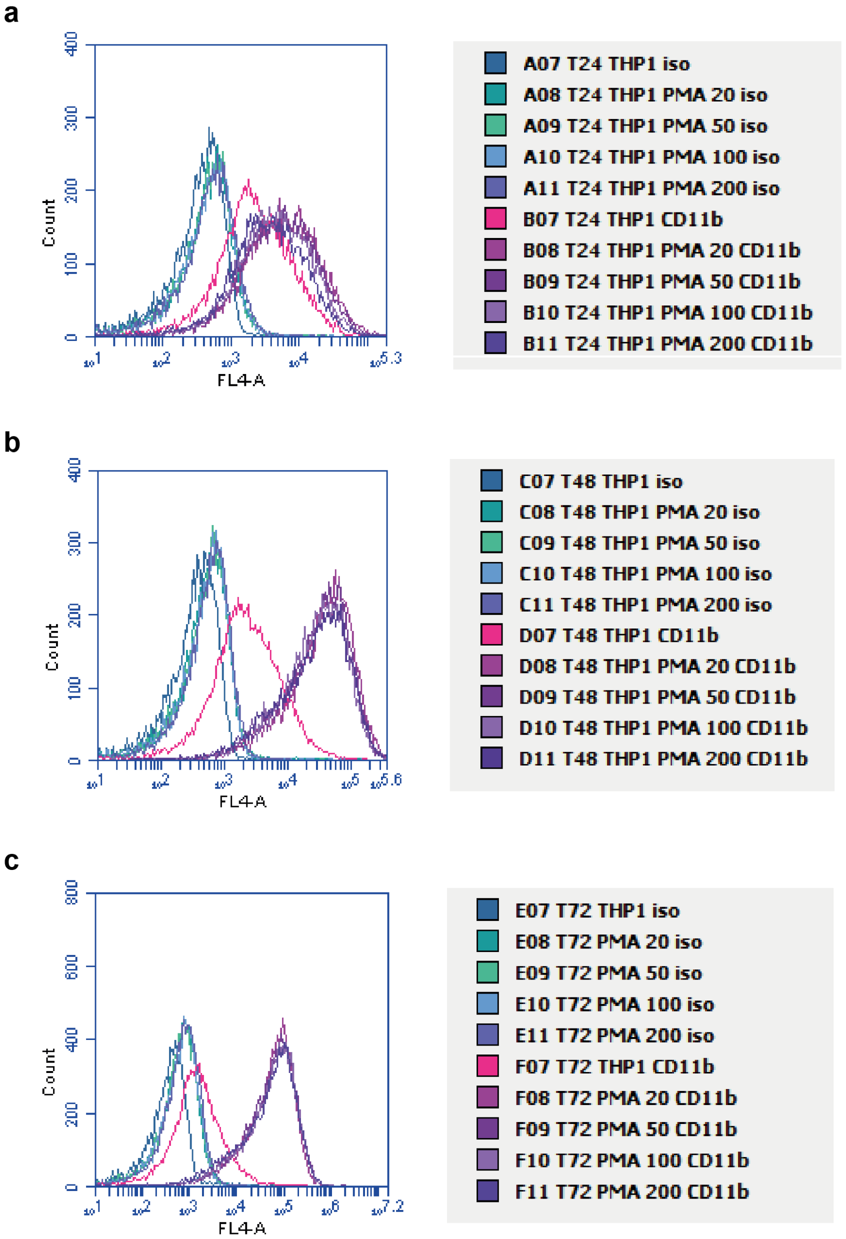


Supplementary Figure 2. Flow cytometry fluorescence intensity (F.I) for CD11b. a. F.I. for THP-1 treated with 20, 50, 100, and 200 nM PMA at 24 hours. b. F.I. for THP-1 treated with 20, 50, 100, and 200 nM PMA at 48 hours. c. F.I. for THP-1 treated with 20, 50, 100, and 200 nM PMA at 72 hours.

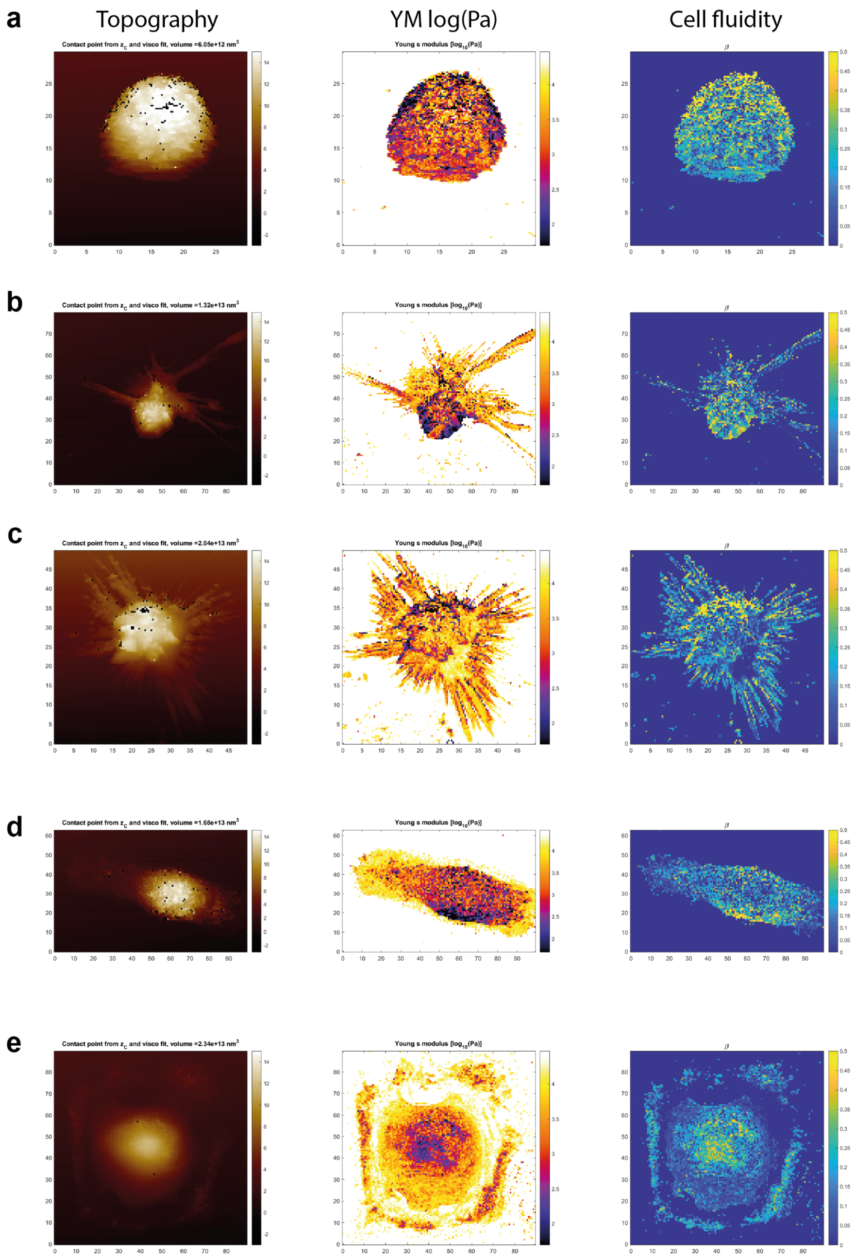


Supplementary Figure 3 a-e. Mechanical maps of THP-1 cells treated with 200nM for 5 days. New phenotypes are observed if the PMA concentration and time of treatment are increased. While we still find round and spread macrophage phenotypes, some stellate cells appear with very long protrusions or filopodia, similar to dendritic cells (b-c). Also, more macrophages migrate in a fused form (d).

Monocytes viscoelastic maps

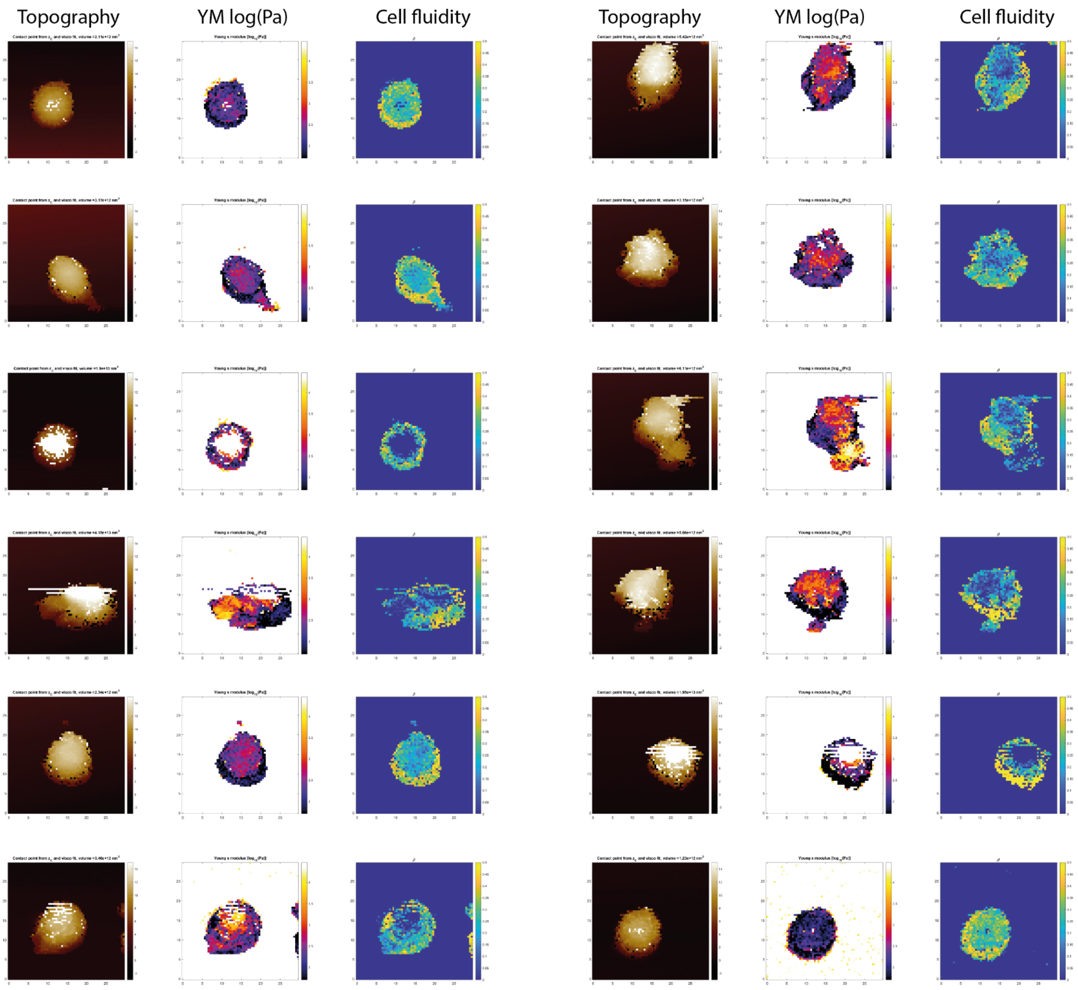

### Round macrophages viscoelastic maps

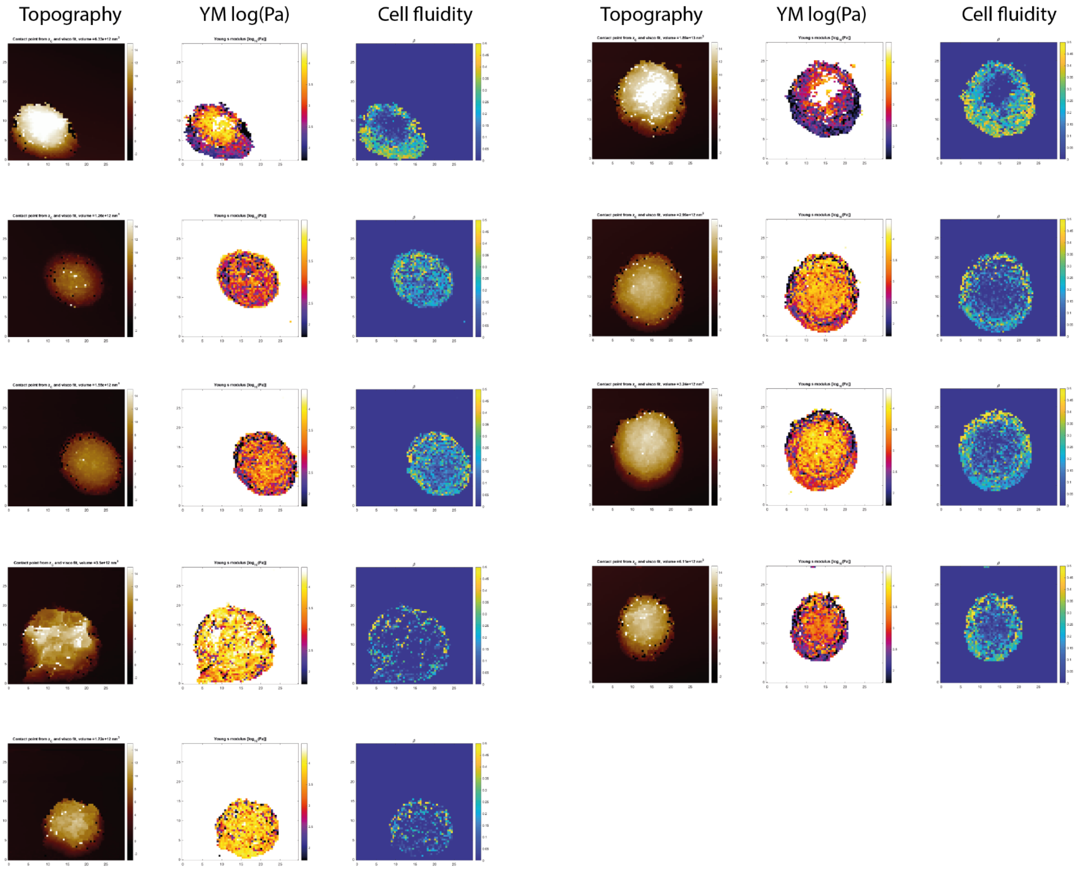


### Spread macrophages viscoelastic maps

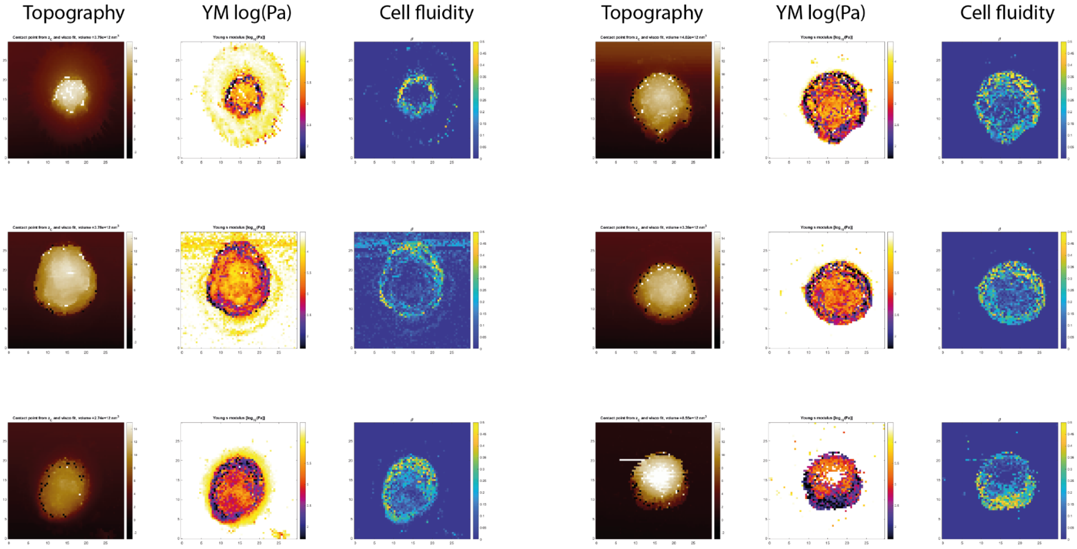

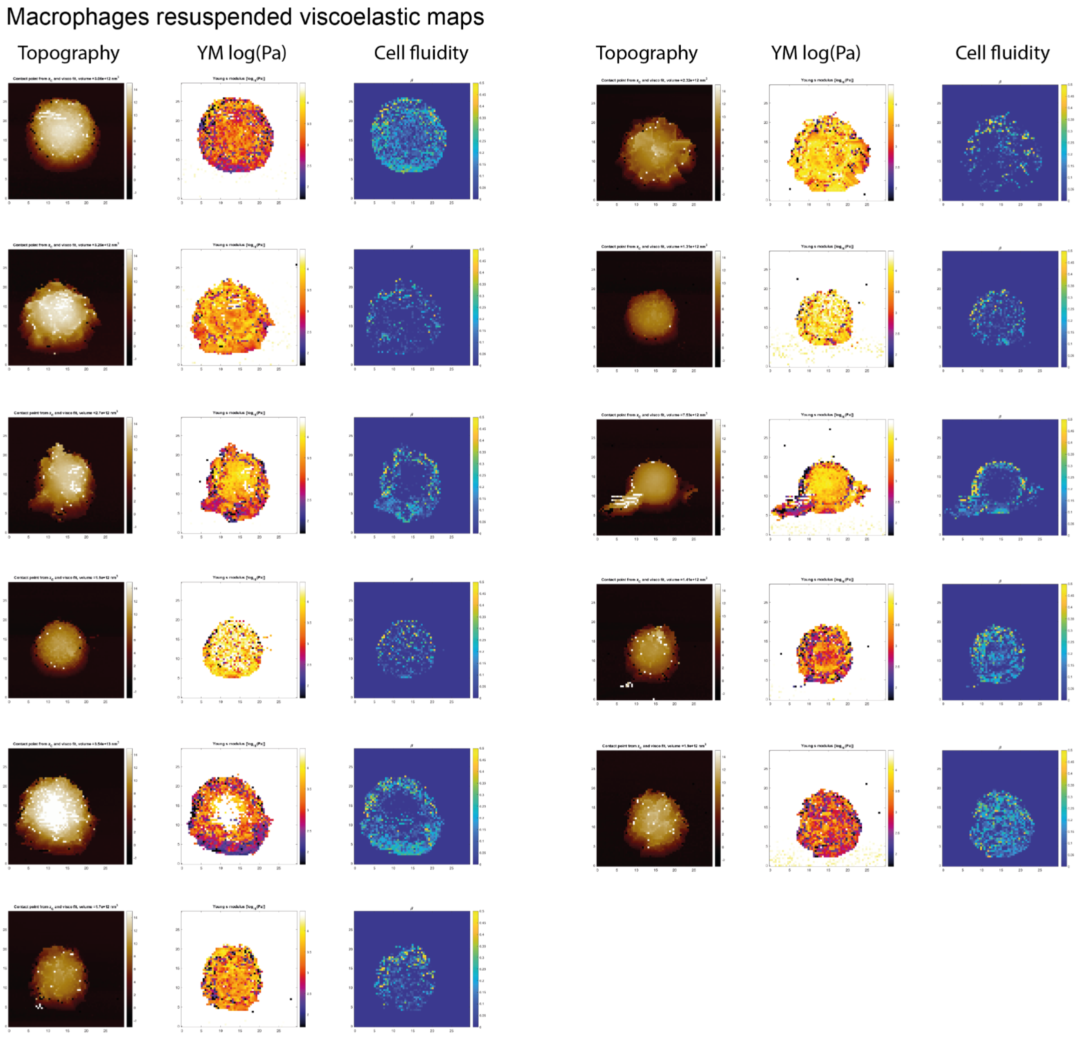


Supplementary Figure 4. Plots single cell elasticity and fluidity maps. All the cell maps that were used for the Figure 3 analysis.

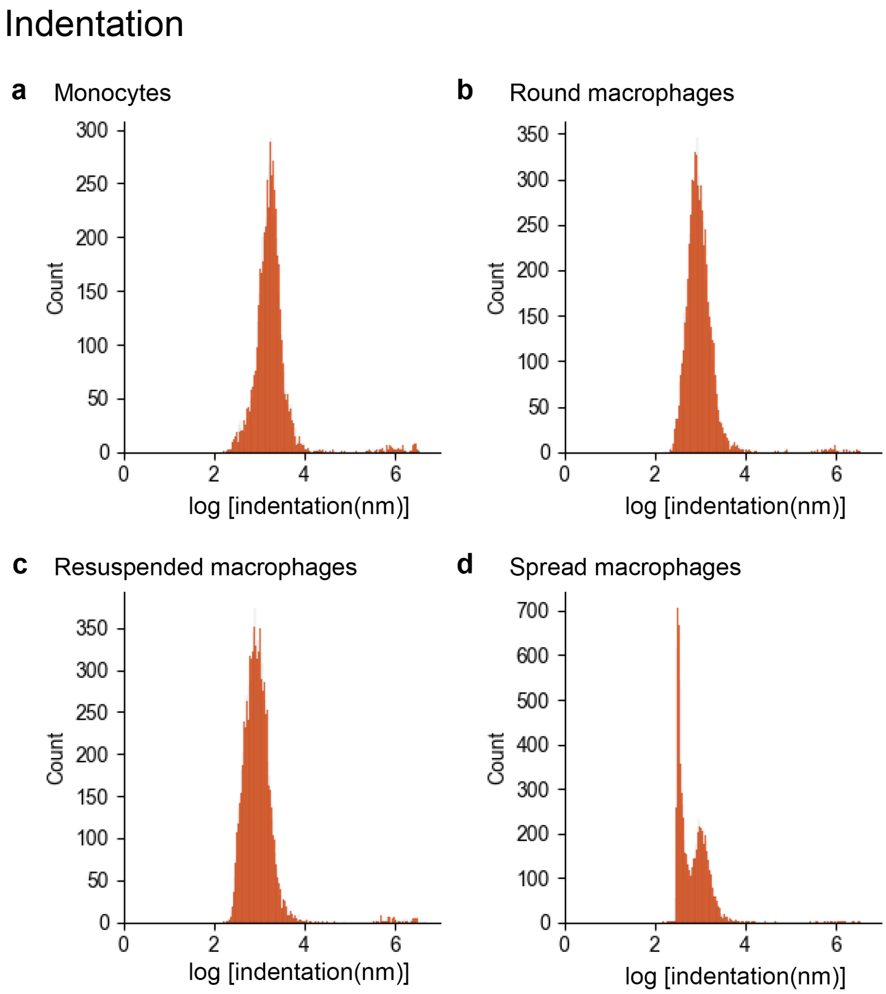


Supplementary Figure 5. Maximum indentation histograms of the studied phenotypes.

Monocytes ICM

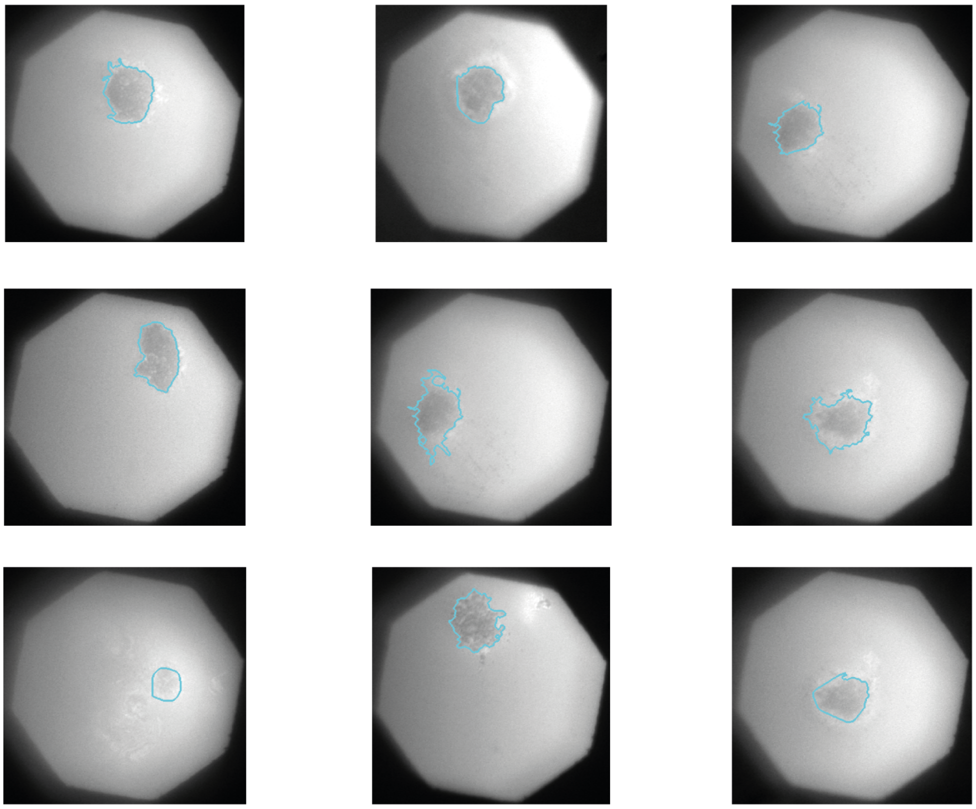

Spread macrophages ICM

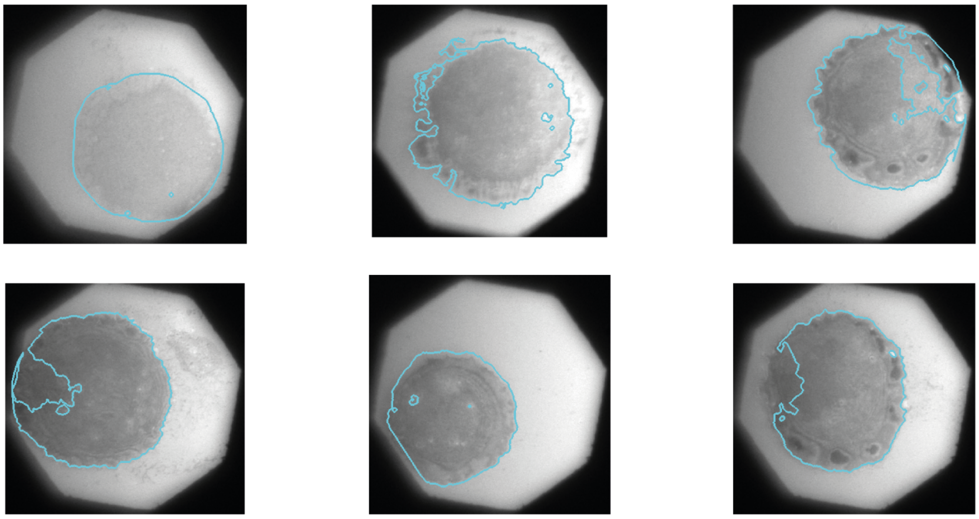

Round macrophages ICM

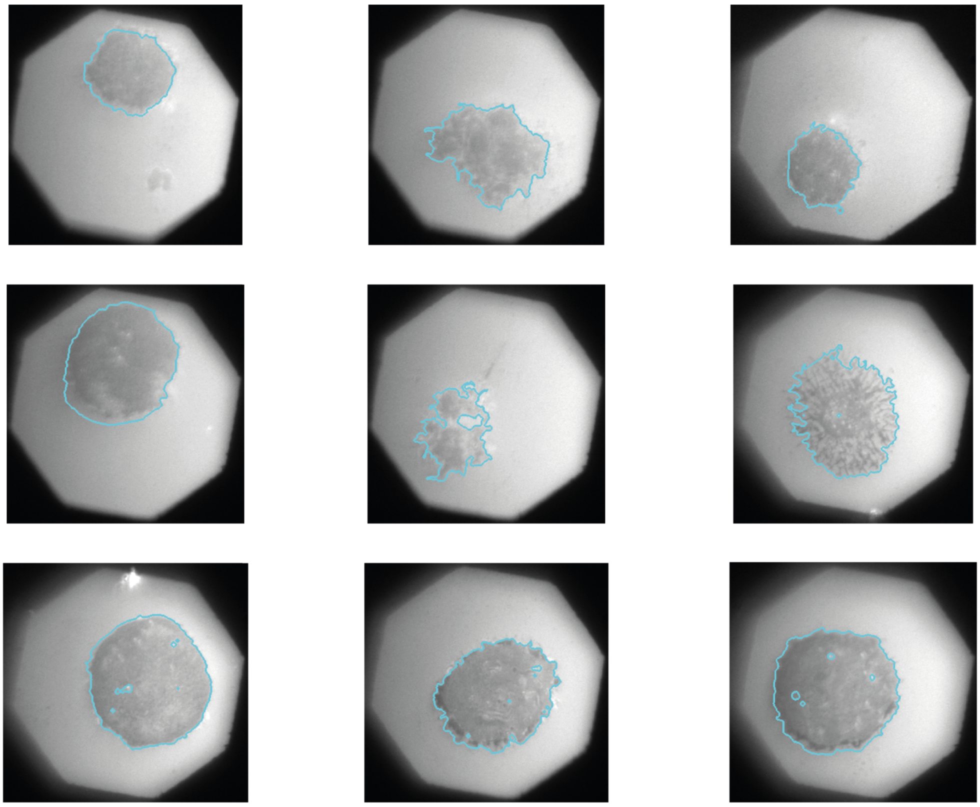

Resuspended macrophages ICM

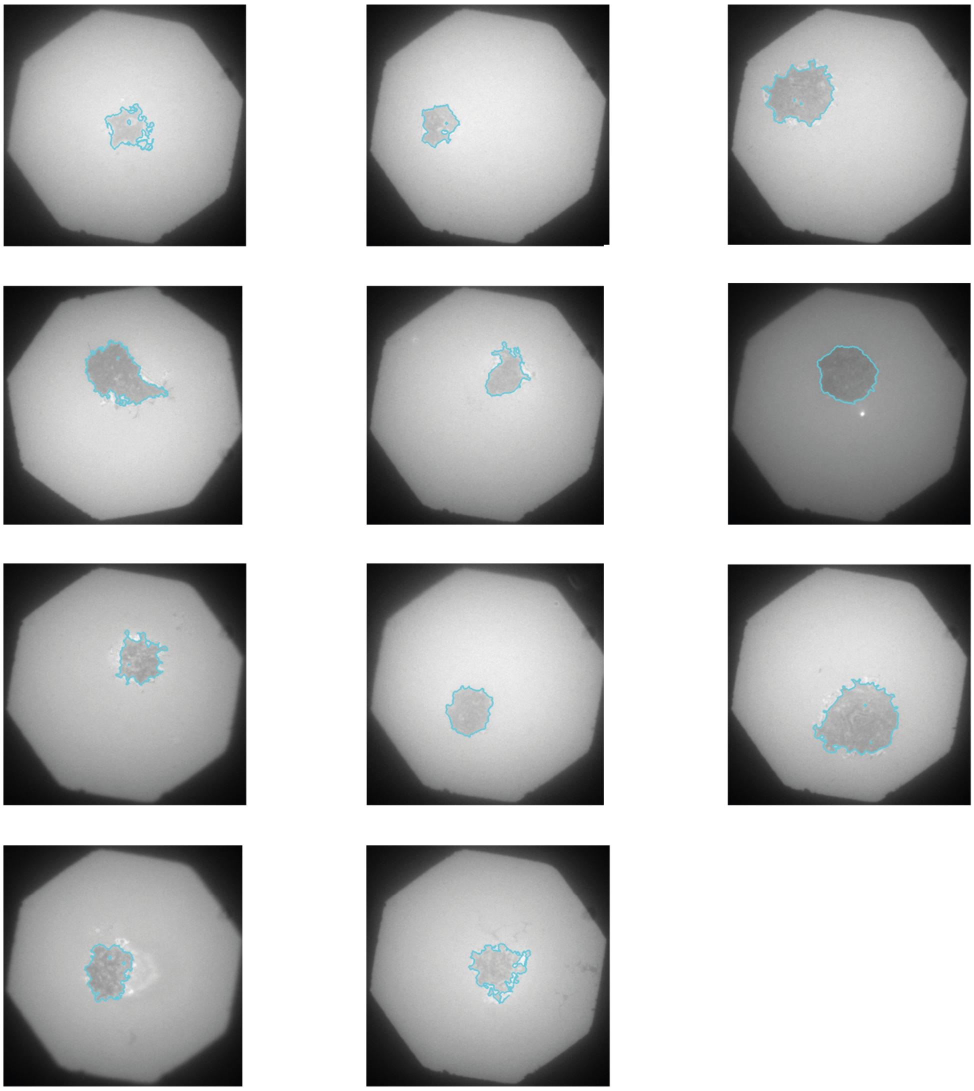

Supplementary Figure 6. ICM single cells images. All the ICM images used in the Figure 4 analysis.

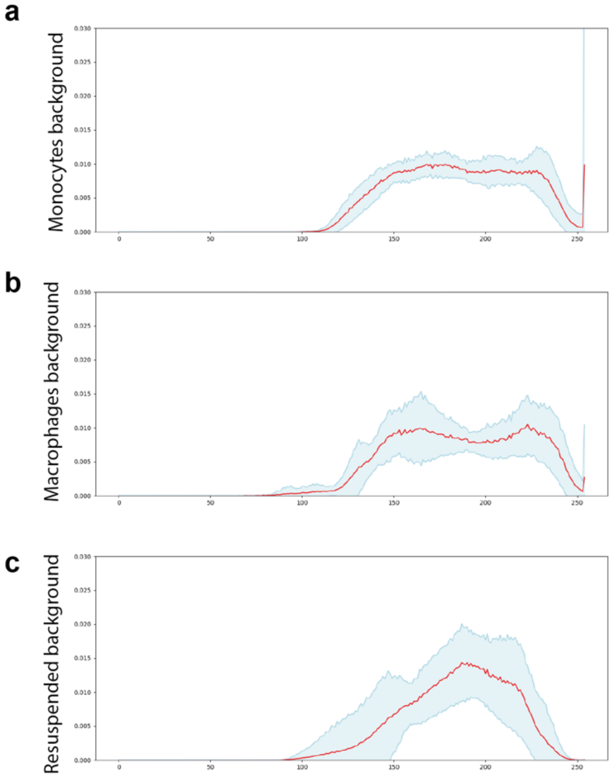


Supplementary Figure 7. a. ICM background intensity for the plates with THP-1 cells (monocytes). b. ICM background intensity for the plates with THP-1 cells treated 48 hours with 20nM PMA (spread and round macrophages). c. ICM background intensity for the plates with THP-1 cells treated 48 hours with 20nM PMA, resuspended with trypsin and immobilized with PLL (resuspended macrophages).
